# NLTD 2.0: A Nonlinear Framework for Robust and Customizable Color Deconvolution in Histopathology

**DOI:** 10.1101/2025.10.02.679756

**Authors:** Florin Selaru, Jude M. Phillip, Denis Wirtz, Pei-Hsun Wu

## Abstract

Advancements in computational approaches have enabled robust utilization of histological tissue data. A crucial step in the development of computational tools for the objective and quantitative analysis of tissue sections has been color deconvolution. Color deconvolution functions by separating the absorption of colors corresponding to stained molecular or tissue compartments. The most widely used color deconvolution method in digital pathology, linear color deconvolution as described in Ruifrok 2001 et al., decomposes color images according to the absorbance values for individual stains. However, linear color deconvolution assumes that stains are linearly decomposable, and it relies heavily upon identifying optimal color vectors of stains, which is often challenging. Furthermore, linear deconvolution methods cannot deconvolve the image with more than three stains, further limiting their broader applicability. To combat the limitations of previous methods, we developed an intuitive and robust color deconvolution method that effectively and accurately separates more than three stain signals, does not rely on predetermined color vectors, and doesn’t rely on identifying optimal stain vectors. The proposed method, NLTD 2.0, presents a robust and efficient solution to tackle color variations in histopathology images, enhancing the reliability and precision of computational pathology. Additionally, incorporating the method as an ImageJ plugin amplifies accessibility and usability, enabling researchers and pathologists to leverage its capabilities without specialized programming skills. The intuitive interface streamlines the application, fostering broader acceptance within the computational pathology community.

## Introduction

Advances in imaging instrumentation and data management have laid the groundwork for computational approaches to analyze digitized images of tissue sections, providing objective, quantitative measurements at the tissue, cellular, subcellular, and molecular levels^1^. Computational pathology offers a cost-effective platform to enhance the throughput, accuracy, and reliability of tissue sample diagnoses^2^. Additionally, the quantitative nature of computational pathology allows for seamless integration with other clinical workflows to enrich pathologists’ understanding of disease, inform treatment strategies, and further stratify patient prognosis. Integrating information from computational pathology with a patient’s clinical data has been shown to produce better prognostic models for many diseases, including prostate cancer^3–5^, lung cancer^6^, breast cancer ^7–11^, glioblastoma^12,13^, basal cell carcinoma^14,15^, and ovarian cancer^16^.

Various studies have established a strong association between tissue and cellular morphology and disease progression as well as survival outcomes^7^.Given the variability in tissue image coloration across batches and institutions, color deconvolution and normalization are critical steps in developing robust prognostic models from histopathological images using modern machine learning and deep learning techniques. For instance, recent research has shown that irregular nuclear morphology in colorectal cancer histology correlates with reduced patient survival^7^, and color normalization enabled Zheng et al. to train a convolutional neural network (CNN) that accurately predicts glioblastoma transcriptional subtypes from histology^17^.

One of the central challenges in computational pathology is the variability in the color appearance of tissue section images across different research laboratories and medical facilities. These variations arise due to differences in tissue fixation, staining protocols, and imaging instrumentation^18^. Previous studies have indicated that even technician variance, and consequently technique differences, can lead to significant differences in stain appearance^19^. Moreover, modifications to conventional hematoxylin and eosin (H&E) staining techniques to reduce material use and processing time^20^ or to enhance contrast and detail in the digital image^21^ have introduced further variability in stain appearance. While these adaptations help pathologists visually, they hamper algorithms that must isolate individual stains (e.g., hematoxylin-labeled nuclei) for downstream analysis.

Several computational approaches, including color deconvolution^22^, histogram equalization^14^, and the use of the CMYK space^23^, have been developed to normalize stain appearance and facilitate tissue type separation^19,24^. Among these, color deconvolution is the most widely used method to extract nuclear and cellular images in both (H&E) and immunohistochemically (3,3’ Diaminobenzidine, DAB) stained images^8,11,22,25^. However, a significant drawback of color deconvolution is its reliance on prior knowledge of each dye’s color spectrum to accurately visualize tissue components^26^. Furthermore, color deconvolution assumes linearity of the color mixing process, which fails to account for non-linear interactions between stains or scanner-specific optical effects^27^.

Due to differences in color appearance between images, using the same stain vector across multiple different images can introduce variance in the deconvoluted image for each dye. Although there are automated methods to determine the optimal stain vector for individual images^28,29^, this additional processing step significantly increases the computational overhead for large image datasets. Furthermore, color deconvolution can separate no more than three colors in a single image, limiting its utility in multiplex immunohistochemistry (mIHC) workflows, where simultaneous detection of four or more biomarkers is increasingly common^30^.

In this work, we present an advanced version of our non-linear tissue-component discrimination method, NLTD 2.0, which significantly enhances the process of registering the color space of histopathology images. NLTD 2.0 extracts color components using a 2D histogram-based approach with tunable parameters to optimize deconvolution outcomes. We demonstrate the method’s robustness across diverse staining protocols and demonstrate its ability to extract more than three color channels, addressing key limitations of linear color deconvolution (LCD).

## Methods

### Images sources

H&E images used in the study were downloaded from Alsubaie et. al. ^31^, Wienert et. al. ^32^, and Mahbod A et. al. ^33^; Immunohistochemically (DAB) stained for LINE-1 ORF1p on ovarian tissue was from ^34^, PTEN DISH assay of prostate cancer tissue was from ^35^, and IHC images are from ^36^; Masson’s Trichrome stained mouse liver tissue was from ^37^ and from mouse tumor tissues generated in-house and stained using standard Trichrome staining protocol that involves deparaffinization, washing, and immersion in trichrome stain solutions to denote cell nuclei, cytoplasm, and collagen fibers.; and pap smear images were downloaded from^38^. The multiplex IHC image in this manuscript were provided by Network for Pancreatic Organ Donors with Diabetes (nPOD) online pathology site.

### Color Deconvolution with NTLD2

In an 8-bit RGB tissue image, the color of each pixel is expressed as a combination of three intensities, red, green, and blue, each ranging from 0 to 255, i.e.,

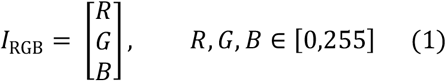

The Optical density (OD) for each pixel was computed using

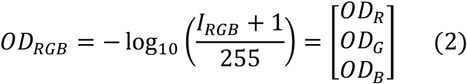

Spherical coordinates representation of optical density was then calculated, to obtain theta (θ), phi (ϕ), and radius (*ρ*), where

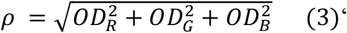

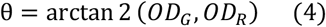

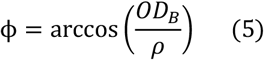

Each pixel in the image was converted from RGB to OD, and then to spherical coordinates (*ρ*, θ, ϕ) to achieve better separation between stains in the color spectrum. Theta (θ), phi (ϕ), and rho (*ρ*) coordinates roughly represent hue, saturation, and absorbance. The theta-phi (θ−ϕ) joint occurrence map (TPOM) of image *I* with *N* pixels for each pixel *i* ∈ {1, 2, …, *N*} is then calculated to represent the 2D color spectrum of the image, i.e.

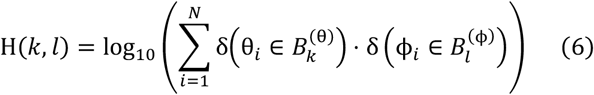

The TPOM has 256 equally spaced bins in both the theta 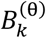, and phi 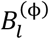. It should be noted that a logarithm scale is applied to the TPOM to improve visual representation of smaller stain populations. In this context, an “occurrence” is simply the number of pixels whose θ and ϕ values fall simultaneously into the *k*_*th*_ θ bin and the *l*_*th*_ ϕ bin. The summation in (6) therefore counts how many pixels share that specific color pair, building up a two-dimensional histogram of joint occurrences. Each delta function, 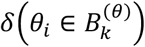 and 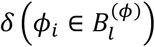 return 1 when the stated condition is true and 0 when it is false, thereby “switching on” the contribution of pixel *i* only when it lies in the specified bin. Multiplying the two delta functions ensures that a pixel is counted in the sum only when both conditions are satisfied.

The segmentation is initialized by enclosing the region corresponding to the highest intensity for each stain in a ROI. Then, the decay parameter (σ) and background cutoff distance are selected to account for the regions of stain mixing and background. All pixel weights are between 0 and 1. Moreover, any bin inside of the ROI is given a weight of 1, while any bins located at a distance greater than the background cutoff value from the ROI are assigned a weight of 0. Any bins between the boundary of the ROI and at a distance less than background cutoff value from the ROI is assigned a weight *w*. This weight *w* is calculated as a function of distance *d* from the ROI boundary and decay parameter σ, defined as follows:

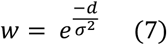

A grayscale image of the same dimensions as the tissue image is then generated, where each pixel is assigned a value based on the designated weight *w* corresponding to the (θ, ϕ) value at the same location in the original tissue image. These weights are then multiplied by the *ρ* value at each corresponding coordinate in the original tissue image. At this point we have a grayscale mask showing the intensity of the stain we deconvoluted in every pixel in the image. Next, we want to convert this back to an RGB 3D image showcasing the deconvoluted stain. To this end, the resulting deconvoluted *ρ* channel produced in the previous step is stacked with the original θ and ϕ channels to create a 3D image. This 3D image is then converted back to cartesian coordinates, creating an OD representation of the deconvoluted image. Finally, the OD image is converted back to the RGB-space, yielding the deconvoluted RGB image for the current stain.

### Immunohistochemistry (IHC) Quantification

A tissue microarray (TMA) of ovarian-cancer tissue stained with an antibody for LINE-1^34^ ORF1 was analyzed with a streamlined optical-density (OD) based approach to benchmark automated IHC scoring. To compare automated scores with the pathologist’s ordinal grades (0 = negative, 3 = strong), each core’s calculated intensity value was plotted against its grade, and the Pearson correlation coefficient was calculated. The resulting correlation (reported as *P*) provided a quantitative measure of concordance, with higher *P* values indicating stronger agreement between the simple-OD based metric and manual immunohistochemistry scoring. Each core was color-deconvoluted by NLTD 2.0 to yield a single image containing only the antigen chromogen. For the analysis described here, we used the deconvoluted *ρ* channel described above rather than the typical cartesian RGB output. Moreover, for consistency the same theta-phi joint histogram ROI and decay value were used to deconvolute all images across all intensity scores. Pre-processing then focused on isolating true chromogen-positive tissue while excluding blank glass and noise: pixels values with *ρ* > *τ*(*τ* = 0.01) were provisionally accepted, after which connected-component filtering removed objects smaller than three pixels. The remaining set of pixels S in the binary mask therefore comprises contiguous regions of isolated antigen staining. Finally, an intensity score was calculated as the mean of the intensity values of all the pixels in *S*. Only chromogen-bearing tissue contributed to *S*. Additionally, unstained areas and scanner artifacts were excluded by the threshold-plus-morphology pipeline, yielding a single continuous value proportional to average antigen load for each TMA core.

### Statistics

Pearson’s product-moment correlation coefficient (*P*) was calculated to quantify the linear relationship between the automated optical-density intensity scores and the pathologist’s ordinal grades.

### Software

All image processing was performed using ImageJ Fiji^39^. All plots were generated using MATLAB 2024a (MathWorks).

### Code availability

The source code is publicly available on GitHub at: https://github.com/DeepBioVision/NLTD-2.0_PlugIn

## Results

### Non-linear Tissue-component Discrimination 2.0 (NLTD 2.0) Method Overview

The NLTD 2.0 method highlights the use of the theta-phi joint occurrence map (TPOM) to separate the chromogen staining of the histology images. The RGB color space can be represented by spherical coordinates (theta, phi, and rho)^40^. In the spherical color space, all normalized color components are represented only in 2-dimensional variables theta and phi. Since the TPOM only considers the normalized color vectors, chromogens with distinct color patterns (such as Hematoxylin vs Eosin) reside at distinct locations on the TPOM map, enabling effective separation (**Fig. 1 a-b**). In this way, the complexity of considering color three dimensionally is circumvented. Compared to the original NLTD^41^ that used the red and blue intensity of pixels to separate the distinct chromogen staining, the TPOM axes provide more orthogonal color information for chromogen staining separation (**Fig. 1 c-d and Supplementary Fig. 1**). The overall NLTD 2.0 method consists of three main steps, detailed in (**Fig. 1e**), including 1) calculating the TPOM map from the input image; 2) selecting the ROIs representing different chromogens in the TPOM maps; 3) creating the color transform function (CTF) based on the ROIs to extract individual chromogen staining from original images (see more details in **Methods**). The deconvoluted tissue component images can be used for further tissue analysis and processing. Additionally, we demonstrate the TPOM map can effectively separate the stains well from H&E images with a variety of color appearances (**Fig. 3**)

**Figure 1.**
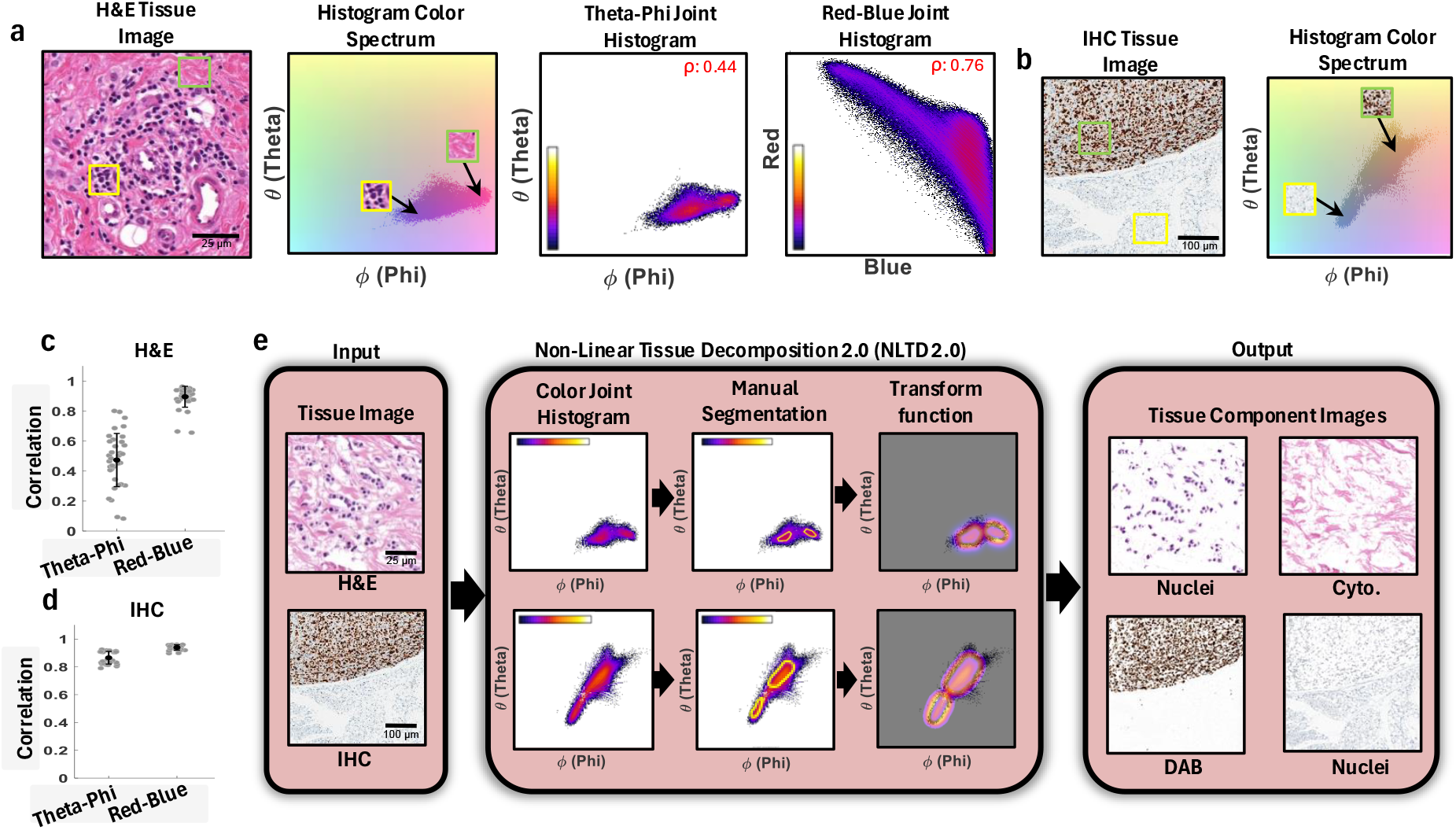
Concept and workflow of NLTD 2.0. **a**. H&E example. Left: raw hematoxylin-and-eosin tile with nuclei (yellow box) and cytoplasm (green box). Second panel: full θ–ϕ color spectrum; boxed regions locate the two stains. Third panel: θ–ϕ joint-occurrence histogram shows two compact clusters (Spearman *ρ* = 0.44). Fourth panel: traditional red-blue histogram yields a streaked distribution (*ρ* = 0.76), indicating stronger channel correlation and poorer separability. **b**. IHC example. An immuno-DAB section (brown chromogen) with counter-stained nuclei is mapped into θ–ϕ space, again revealing distinct clusters for each stain. **c-d**. Quantitative cluster separability. For 15 heterogeneous H&E images (c) and 10 IHC images (d), the θ–ϕ histogram consistently exhibits lower intra-cluster grayscale correlation than the red-blue histogram (mean ± SD shown), confirming improved orthogonality of the spherical representation. **e**. NLTD 2.0 pipeline. Starting from an input bright-field image (left column), NLTD 2.0 builds a θ–ϕ joint histogram, the user (or algorithm) selects stain-specific regions, and a decay-weighted transform function is generated. Applying this function produces deconvoluted tissue-component images (right column), e.g., nuclei vs. cytoplasm for H&E, or DAB vs. hematoxylin for IHC.

### NLTD 2.0 For Robust Customizable Color Deconvolution

Identifying the ROIs corresponding to different staining on the TPOM is key to NLTD 2.0. In most H&Es (Hematoxylin and Eosin images), the stained colors are readily distinguishable in the TPOM (**Fig. 1e**). However, in certain cases, the stain locations on the TPOM might be more overlapping and less straightforward to determine. In this case, examining the TPOM at a specific rho range can aid the selection of stain ROIs since the visible difference between stains is associated more with rho when it is less associated with theta and phi (**Fig. 2a and Supplementary Fig. 2**). As a note, absorbance is a proxy for rho that is easier to understand. In the example image, we show different tissue components are also visually distinguishable from the absorbance (rho) image (**Fig. 2a**). Histogram analysis of the absorbance of the H&E image used shows three distinct absorbance subpopulations that were found corresponding to eosin, hematoxylin, and blood, which ranked from low absorbance to medium to high absorbance, respectively (**Fig. 2b**). Therefore, examining the TPOM at various ranges of absorbance can further improve the identification and the ROI corresponding to individual staining/color components.

**Figure 2.**
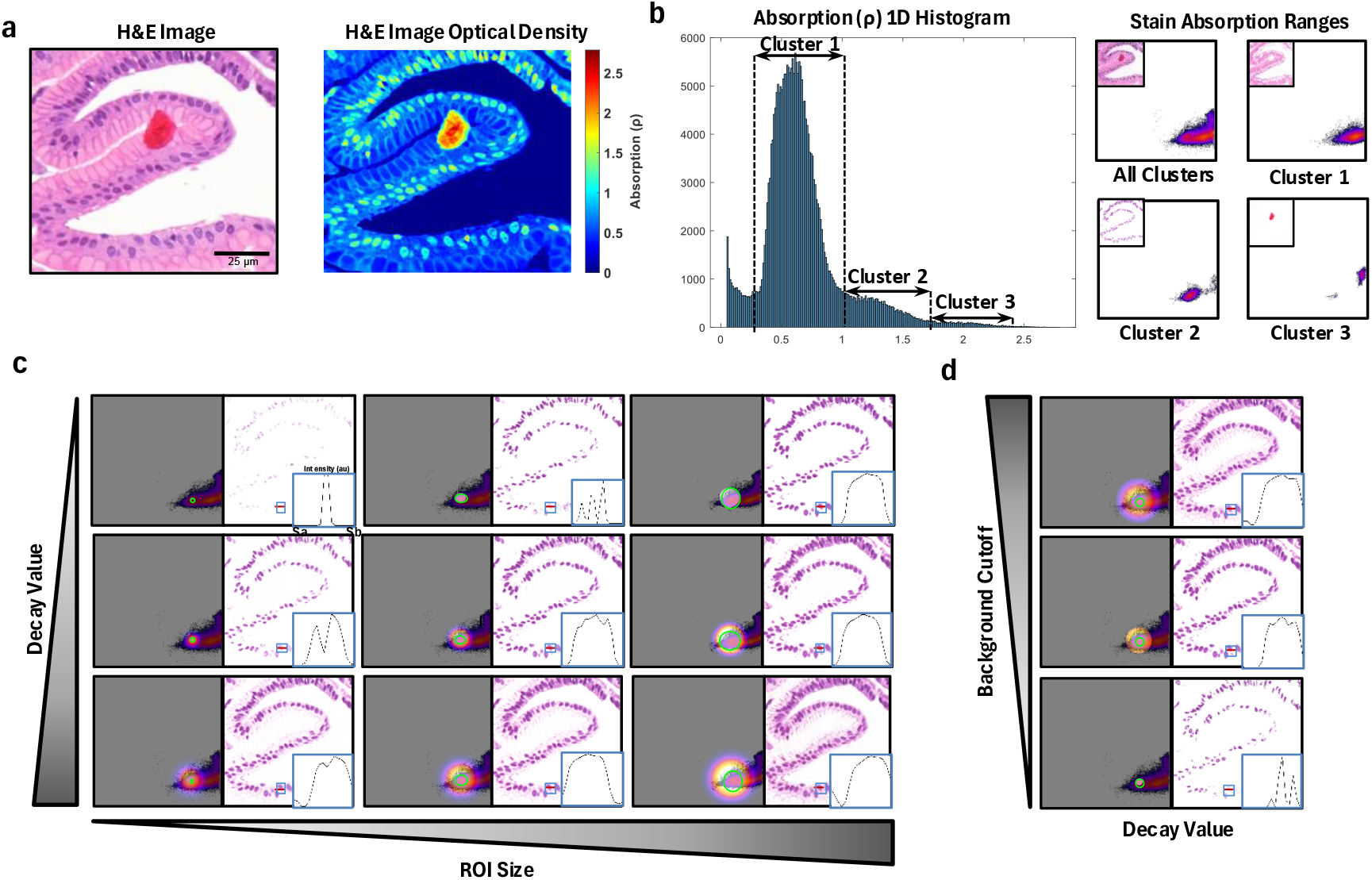
Tunable parameters of NLTD 2.0 enable precise stain isolation. **a**. Absorbance accentuates chromogen differences. Left: an H&E tile containing cytoplasm (eosin), nuclei (hematoxylin), and erythrocytes. Right: the same tile rendered as optical-density (*ρ*) heat-map; higher absorbance (warm colors) highlights densely stained nuclei and blood. **b**. One-dimensional *ρ* histogram separates three stain populations. The histogram reveals three absorbance clusters (dashed boxes): Cluster 1 = eosin-rich cytoplasm, Cluster 2 = hematoxylin nuclei, Cluster 3 = blood. TPOM sub-panels (right) show how restricting *ρ* to each cluster localizes the corresponding stain in θ–ϕ space and in the image. **c**. Joint effect of ROI size and decay parameter on deconvolution quality. Nine combinations are displayed (ROI diameter decreases left→right; decay increases bottom→top). Each cell shows: left, θ–ϕ histogram with the selected ROI (green); right, resulting nuclei channel with an inset line-profile through the same nucleus (blue box). Large ROIs or high decay include more mixed pixels, softening boundaries; small ROIs or low decay sharpen separation but risk losing faint nuclei signal. **d**. Background-cutoff distance governs suppression of non-target stains. With a fixed decay, raising the cutoff (bottom→top) progressively excludes distant colors, reducing eosin bleed-through in the final nuclei image (right of each pair). Inserts show corresponding intensity profiles.

NLTD 2.0 generates deconvoluted images based on the color transform function (CTF). The CTF deconvolves images by determining which parts of the color spectrum belong to each selected component based on the ROI, decay value, and background cutoff distance. Colors outside the selected ROI have a variable weighting determined by the decay value and background cutoff distance (more details in the **Methods** section).

The CTF is customizable for achieving optimal deconvolution outcomes. A smaller ROI typically associates a smaller proportion of the color spectrum with the stain at the risk of missing parts of the color spectrum at the boundary of the current stain. Conversely, a larger typical ROI associates a larger proportion of the color spectrum with the stain at the risk of including parts of the color spectrum belonging to other stains as well (**Fig. 2c**). Moreover, a bigger decay value results in softer transitions between the parts of the color spectrum included in the stain and excluded from the stain. In contrast, a smaller decay value results in sharper transitions between the parts of the color spectrum included in the stain and excluded from the stain (**Fig. 2c**). Adjusting background cutoff distance can minimize the interference from colors belonging to other stains and reduce background noise by selecting distance from the boundary of the ROI at which the color spectrum becomes fully excluded from the stain (**Fig. 2d**). Hence, the deconvolution performance can be effectively adjusted and optimized in the NLTD 2.0. In comparison, linear color convolution offers no further adjustability after the color vectors are determined, contributing to suboptimal results (**Supplementary Fig. 3**).

### Robustness of NLTD 2.0 in H&E Image Deconvolution

We further demonstrate robust performance of NLTD 2.0 by deconvoluting hematoxylin (i.e. nuclei color component) from a set of H&E images with largely varying color appearance. The TPOM readily distinguishes the distinct color appearance in each H&E image, which is shown by the nuclei color component of each image appearing in a distinct location on the TPOM (**Fig. 3**).

**Figure 3.**
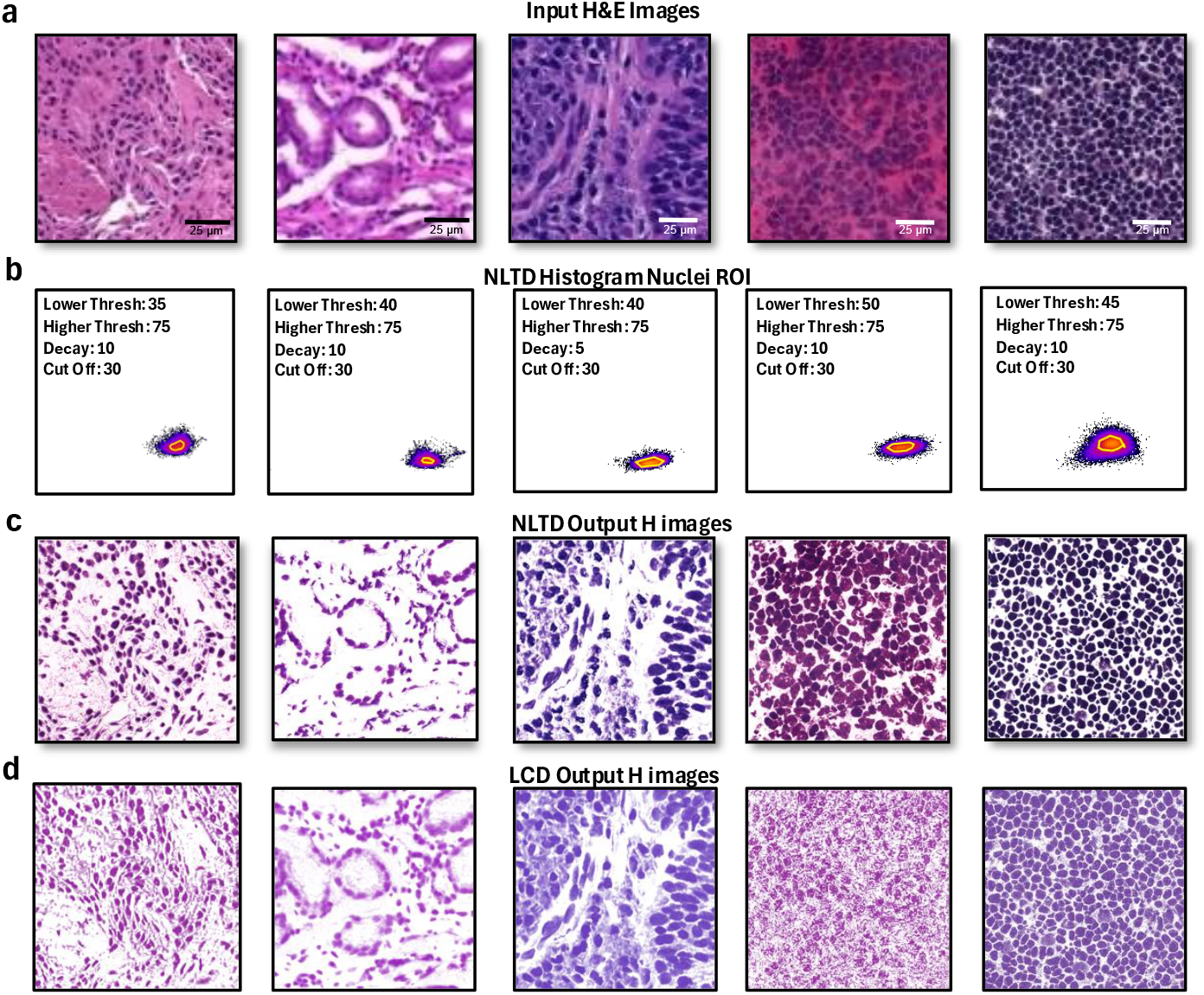
NLTD 2.0 reliably isolates hematoxylin in heterogeneous H&E samples. **a**. Five representative H&E tiles with markedly different stain hues (top row) were processed through NLTD 2.0 and linear color deconvolution (LCD). **b**. θ–ϕ joint-histogram for each tile, with the nuclei region of interest (ROI) highlighted. Text in each panel lists absorbance lower/higher thresholds, decay factor, and background cut -off used for that sample. **c**. Nuclei channels produced by NLTD 2.0 using the corresponding ROI and parameters above. Note the crisp nuclear boundaries and minimal cytoplasmic bleed-through across all color variations. **d**. Nuclei channels obtained with optimized linear color deconvolution. LCD leaves residual eosin signal and blurs nuclear details, especially in tiles with atypical staining. Collectively, the panel demonstrates that NLTD 2.0 maintains consistent, high-quality hematoxylin extraction despite inter-slide stain heterogeneity, outperforming conventional LCD.

We examined the TPOM corresponding to the Hematoxylin by restricting the range of absorption in the TPOM to help facilitate robust identification and selection of Hematoxylin stain TPOM (**Fig. 3**). Notedly, despite the large color differences between the images, similar absorbance ranges were needed to isolate each Hematoxylin signal. In general, we found that a lower threshold of 2 and upper threshold of 3.75 isolated the Hematoxylin signal well (**Fig 3**). Additionally, we found that in general a decay of 10 and background cutoff distance of 30 led to optimal deconvolution results (**Fig 3**). We next compared the performance of the deconvolution with the linear color deconvolution (LCD). Utilizing the preset color vectors^14^ in LCD results in poorly deconvolved images compared to NLTD 2.0 **(Supplementary Fig. 3)**. Even when the color vectors for LCD were optimized, the results remained suboptimal in comparison to NLTD 2.0 **(Fig 3)**. Using LCD with optimized color vectors, the boundaries of nuclei (Hematoxylin) are not as clearly resolved and include substantial signals from non-nuclei stain colors (such as stromal tissue). Overall, these results demonstrate NLTD 2.0’s robust and effective process in deconvolving H&E tissue images, regardless of color differences.

### NLTD 2.0 for quantitative IHC analysis

We next demonstrate that NLTD 2.0 produces quantitative, pathologist-concordant IHC readouts (**Fig. 5**). We applied NLTD 2.0 to an ovarian cancer tissue microarray (TMA) cohort that had been immunolabeled for L1ORF1p, a cytoplasmic protein linked to cancer^42^ to deconvolved the DAB (3,3’-Diaminobenzidine) stains. The DAB-stained level in each tissue core was scored by a trained pathologist on a discrete scale of 0 to 3, where 0 indicates no detectable protein expression and 3 represents high expression. The same parameters in NLTD 2.0 were used for deconvolved DAB-stained images in all tissue cores. The stained level of each tissue core was then scored based on the average absorbance in the deconvolved DAB images in the pixels with colors corresponding to the selected color ROI in TPOM. The DAB scores by NLTD 2.0 are highly associated with the pathologist’s assessment with a Spearman correlation (ρ) of 0.87 (**Fig. 5a**) which is better than previously reported (ρ = 0.81) ^41^ and better than LCD (ρ = 0.67) (**Fig. 5b**). Representative IHC images across grades 0–3 illustrate the expected progression in chromogen signal (**Fig. 5c**). Together, these data show that NLTD 2.0 provides a robust, nonparametric, and quantitative DAB metric that aligns closely with expert assessment while exceeding LCD performance for IHC scoring.

### Implementing NLTD 2.0 in images beyond dual color staining

Current color deconvolution methods, primarily based on linear color deconvolution, have inherent limitations that restrict their application to images containing at most three color components, including the background^40^. In contrast, NLTD 2.0 overcomes this limitation by allowing the selection of any number of color components.

To illustrate this capability, we applied NLTD 2.0 to pancreatic tissue sections stained using multiplex immunohistochemistry (mIHC) with four distinct markers (**Fig. 4a**). We showed that the TPOM distinctly displayed four separate clusters corresponding to each marker, enabling clear identification, and selecting stained ROIs corresponding to each staining component for deconvolution. Compared to linear color deconvolution (**Supplementary Fig. 3**), which failed to adequately separate overlapping stains and resulted in substantial cross-stain contamination, NLTD 2.0 provided significantly cleaner, clearer, and more precise stain separation.

**Figure 4.**
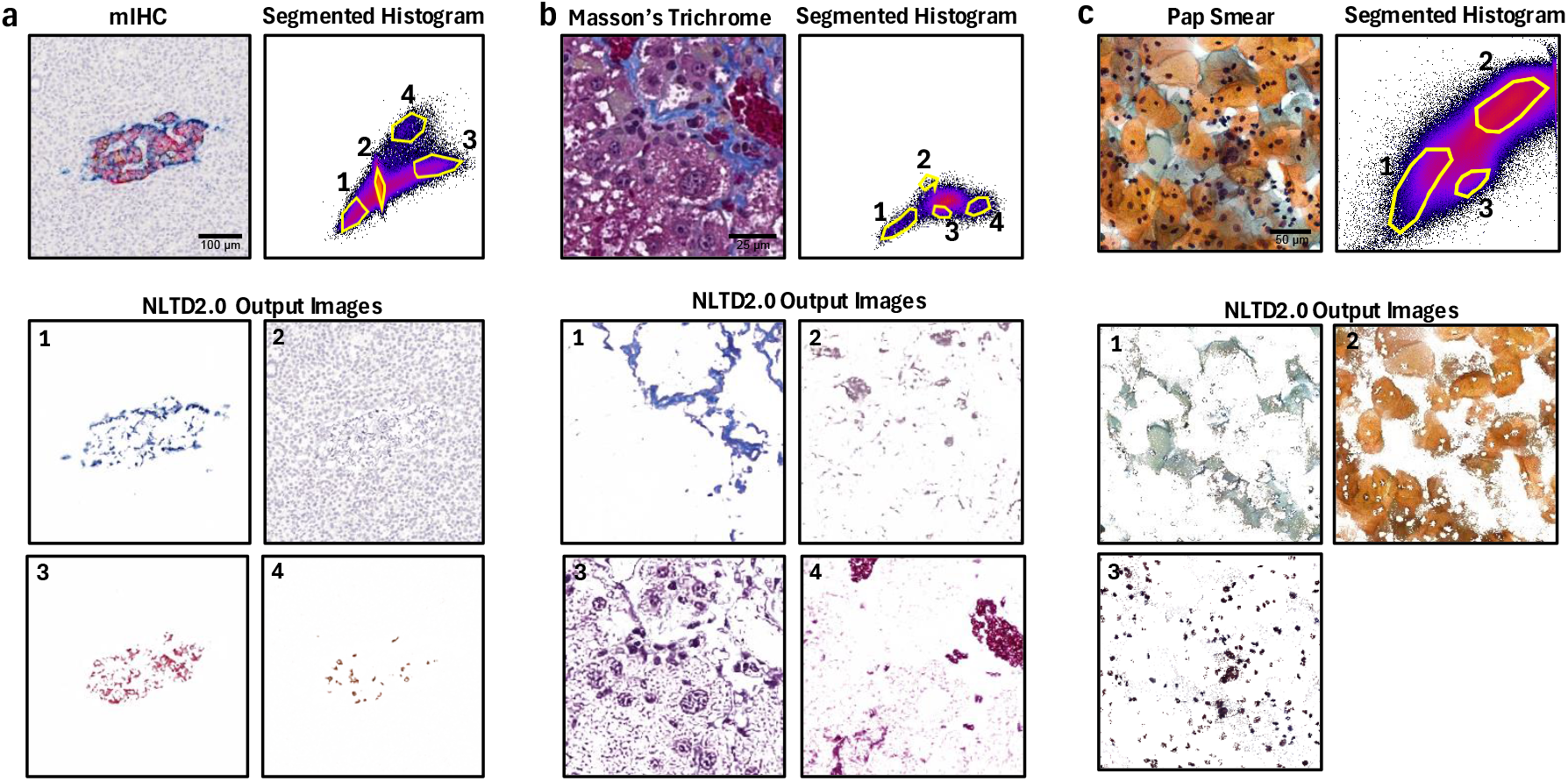
NLTD 2.0 separates four or more chromogens across diverse bright-field preparations. **a**. Four-plex chromogenic IHC. Left: raw multiplex IHC tile. Middle: θ–ϕ histogram with four stain-specific ROIs (yellow, numbered 1-4). Bottom: NLTD 2.0 outputs for each chromogen, showing clean separation of all four labels. **b**. Masson’s Trichrome. Left: original section. Right: TPOM with four ROIs. Bottom: NLTD 2.0 outputs reveal collagen (1), cytoplasm (2), nuclei (3), and erythrocytes (4) as distinct channels. **c**. Conventional Pap smear. Left: Pap cytology image. Middle: histogram with three ROIs capturing nuclei (1), cytoplasm (2), and background debris (3). Bottom: NLTD 2.0 outputs for each component, illustrating accurate extraction even in loosely adherent cellular smears.

**Figure 5.**
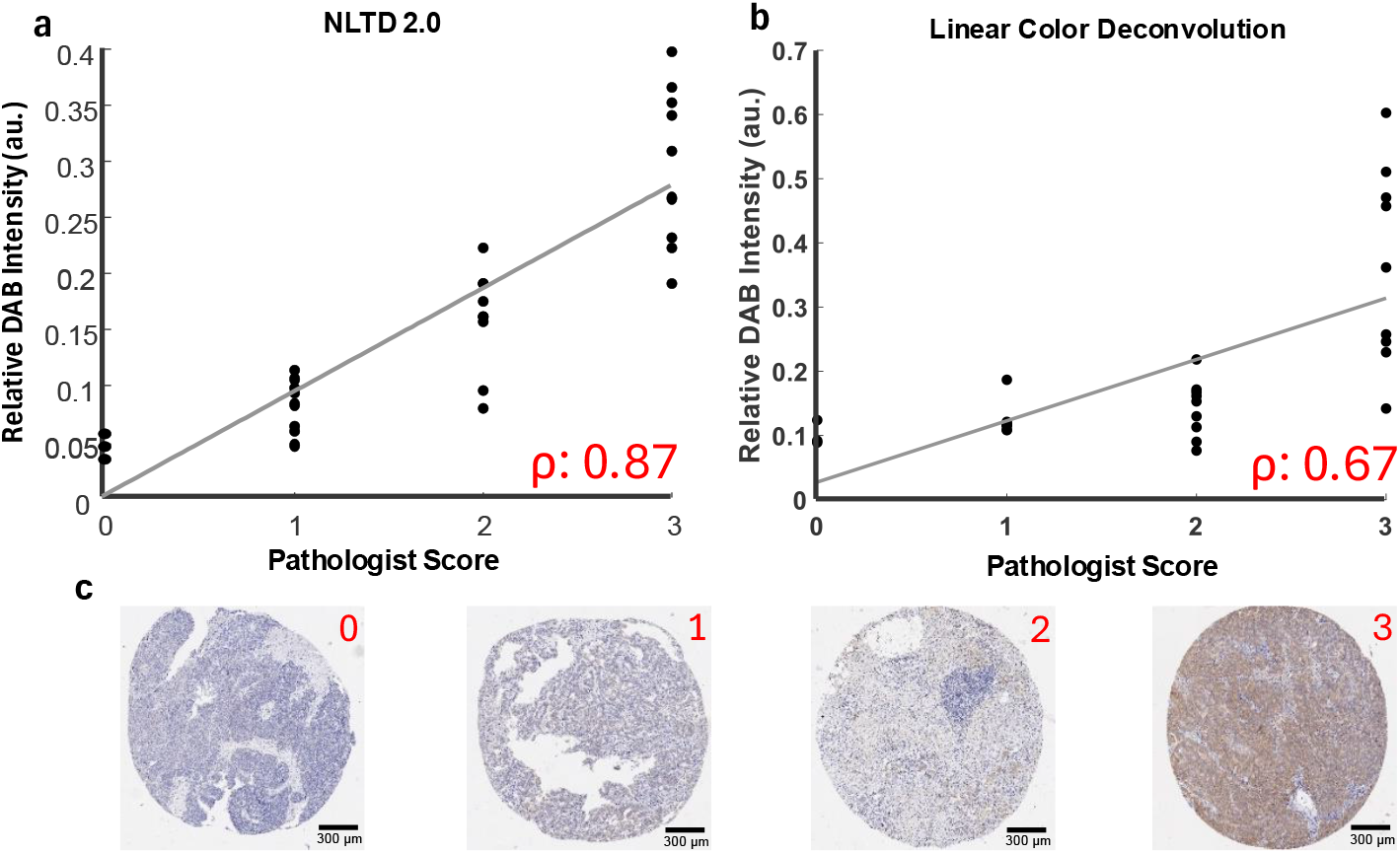
NLTD 2.0 provides quantitative, pathologist-concordant DAB scoring. **a-b**. Scatter plot shows the mean DAB optical-density (y-axis: NLTD 2.0 output (a), Linear Color Deconvolution (b)) versus ordinal pathologist grades (x-axis, 0–3) for an ovarian-cancer tissue-microarray cohort (n = 40 cores). Each dot represents one core; grey line, least-squares fit. The automated intensity metric correlates strongly with manual assessment (Spearman *ρ* = 0.87 for NLTD 2.0 and 0.67 for Linear Color Deconvolution, annotated in red). **c**. Representative deconvoluted cores for grades 0, 1, 2, 3 illustrate the progressive increase in chromogen signal.

Additionally, we demonstrated NLTD 2.0’s robust performance by successfully deconvolving four color components from Masson’s Trichrome-stained tissue sections, clearly separating collagen fibers, muscle tissue, erythrocytes, and cytoplasmic components^43^ (**Fig. 4b**). Furthermore, we applied NLTD 2.0 to Pap smear images, effectively distinguishing cellular structures, nuclei, cytoplasm, and background staining (**Fig. 4c**). These results underscore the ability of NLTD 2.0 to manage stains across diverse histological preparations reliably.

## Discussion

In this work, we established NLTD 2.0 as a robust method for stain separation in histopathology. We demonstrated that NLTD 2.0 effectively produces consistent, quantitative representations of tissue components across images with significant staining variability. By making the workflow accessible via an ImageJ plugin, we promote broad adoption of the method in the computational pathology community.

Various advanced approaches have also been explored to address stain variability, including machine learning algorithms (e.g., deep learning–based stain normalization)^44^, non-negative matrix factorization (NMF) techniques^28^, and singular value decomposition (SVD) based techniques. However, these methods can be computationally intensive (NMF), require extensive training data (deep learning), or are restricted to extracting at most three stain vectors (NMF and SVD), limiting their broad adoption in routine workflows. By comparison, linear color deconvolution (LCD) has historically played a crucial role in enabling quantitative and objective tissue assessment due to its simplicity and effectiveness^40^. However, LCD methods are hindered by stain heterogeneity, a fixed number of color components, and limited adaptability. The method introduced in this study, NLTD 2.0, addresses these limitations by providing a highly flexible and efficient approach to color deconvolution.

Unlike many conventional color deconvolution methods, NLTD 2.0 operates without prior color assumptions, enabling it to consistently process a wide range of images regardless of stain variability or the number of color components (**Fig. 4**). This adaptability makes it particularly useful for analyzing heterogeneous datasets and publicly available histopathology images, reducing the need for strict in-house image preparation. Additionally, NLTD 2.0 leverages a two-dimensional color space, significantly lowering computational costs while maintaining high accuracy. Its four intuitive parameters allow for user-defined adjustments, ensuring reproducibility and customization across different applications (**Fig. 3**).

NLTD 2.0 effectively manages multiplex immunohistochemistry, Masson’s Trichrome, and Pap smear preparations, highlighting its broad applicability. Its implementation as an ImageJ plugin ensures ease of use and widespread accessibility, even for researchers without specialized computational skills. Computational pathology is rapidly transforming diagnostic and prognostic capabilities, providing insights beyond those available through traditional histological examination alone. NLTD 2.0 offers an accessible and robust deconvolution platform that can be used by researchers without specialized expertise in computational analysis or pathology. We have implemented the NLTD 2.0 method as a plug-in on the open-source and widely used ImageJ platform to ensure broad accessibility and to facilitate large-scale histopathological image analysis.

Despite these advantages, the current dependence on manual region selection introduces some variability. Future developments should focus on integrating NLTD 2.0 with automated, objective segmentation methods to further reduce user bias. Additional validation across diverse clinical datasets is also essential to refine its robustness for clinical application. While NLTD 2.0 overcomes many limitations of traditional color deconvolution methods, future work should explore its integration with machine learning–based image segmentation and classification techniques. Additionally, further validation on high-throughput clinical image data will be critical to refining its robustness in real-world applications. By enabling improved tissue analysis with minimal computational overhead and no requirement for prior stain information, NLTD 2.0 has the potential to advance diagnostic and prognostic capabilities in pathology, contributing to better patient outcomes.

## Abbreviations

(H&E): Hematoxylin & Eosin
(DAB): 3,3’-Diaminobenzidine
(IHC): Immunohistochemistry
(mIHC): Multiplex Immunohistochemistry
(NLTD): Non-Linear Tissue-component Discrimination
(LCD): Linear Color Deconvolution
(TPOM): Theta-Phi joint Occurrence Map
(TMA): Tissue Microarray
(OD): Optical Density
(CTF): Color Transform Function
(NMF): Non-Negative Matrix Factorization

## Additional Information

### Author contributions

F.S, P.H.W. and D.W. designed the experiments. F.S, P.H.W., J.P collected the data; P.H.W., and F.S. developed the analytical tools and analyzed the data. F.S., P.H.W., and D.W. wrote and edit manuscript.

## Acknowledgements

The authors acknowledge the following sources of support: UG3CA275681 (PHW), UH3CA275681(PHW); U54AR081774 (DW); U54CA268083 (DW); R01CA300052 (DW); R35GM157099 (JMP), all from the National Institutes of Health. Multiplex immunohistochemistry Images in this manuscript were provided by Network for Pancreatic Organ Donors with Diabetes (nPOD) online pathology site. nPOD is a collaborative type 1 diabetes research project supported by Breakthrough T1D and the Helmsley Charitable Trust. Organ Procurement Organizations partnering with nPOD to provide research resources are listed at https://npod.org/for-partners/npod-partners/

