## Supplementary Figures for "NLTD 2.0: A Nonlinear Framework for Robust and Customizable Color Deconvolution in Histopathology"

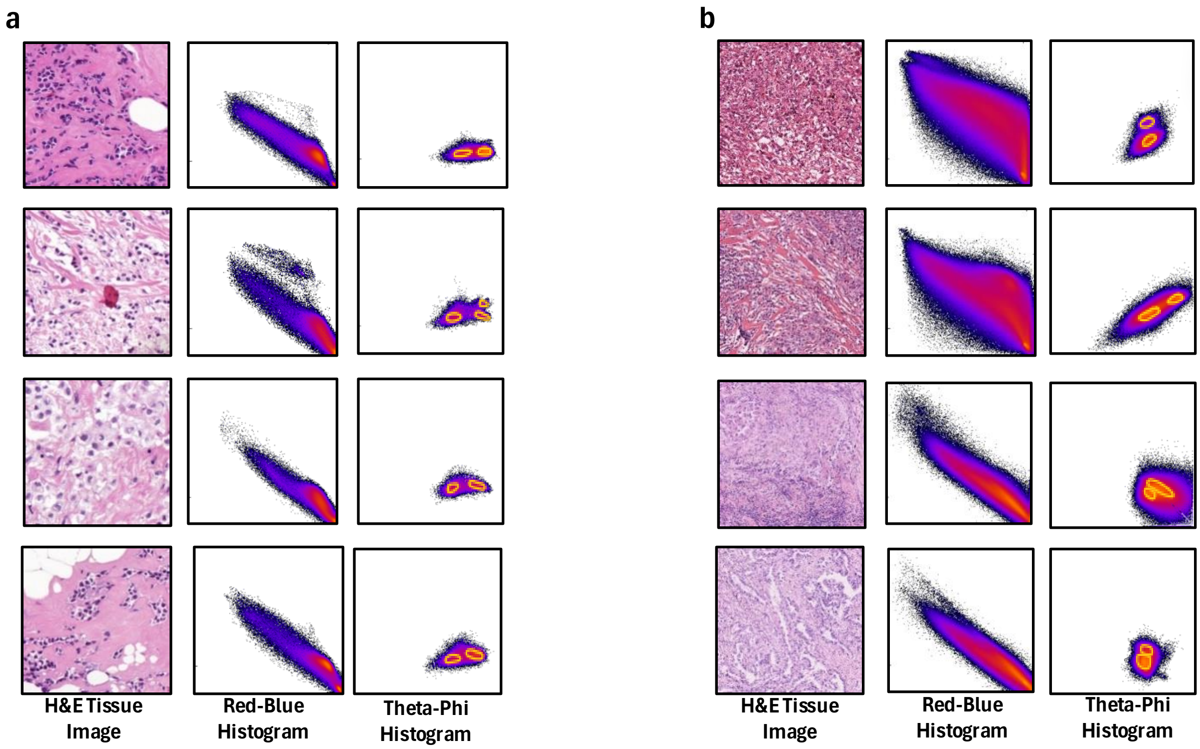


**Supplementary Fig 1. R-B histogram vs Theta-phi histogram**

**a-b.** Both (a) and (b) are in the same format showing H&E tissue images in their left most column, the resulting Red-Blue Histogram for each image in the middle column, and the Theta-Phi Occurrence Map (TPOM) in the right column with the present stain signals (and blood for one image in (a)) circled. This aims to highlight the improved separation in stain signal from the previous method using a Red-Blue Histogram to the TPOM for an assortment of different tissues stained with H&E.

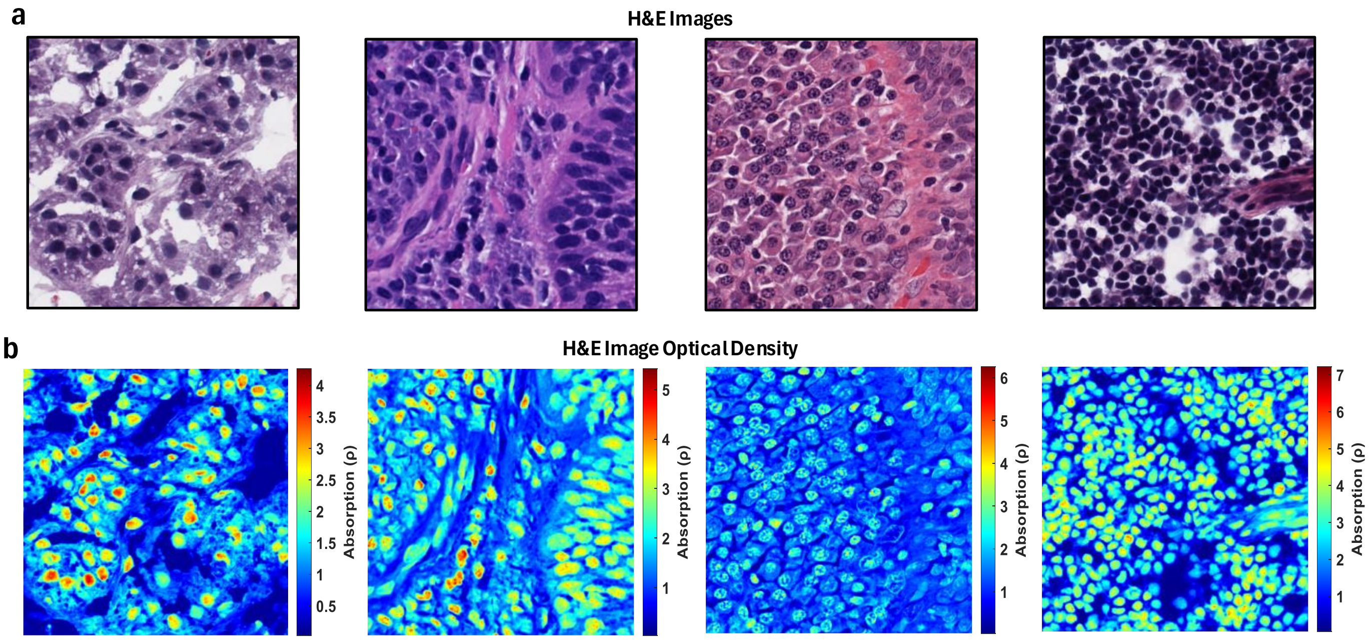


**Supplementary Fig 2. absorbance of H&E images are associated with chromogenic staining.**

**a.** This row displays an assortment of H&E-stained tissue images with great color differences between them, aiming to generalize the observation in this figure. **b.** Each H&E tissue image has its optical density absorbance channel plotted in this row showing that absorbance thresholding can be a useful tool for separating the contributions between Hematoxylin and Eosin, validating the implementation of the absorbance thresholding feature.

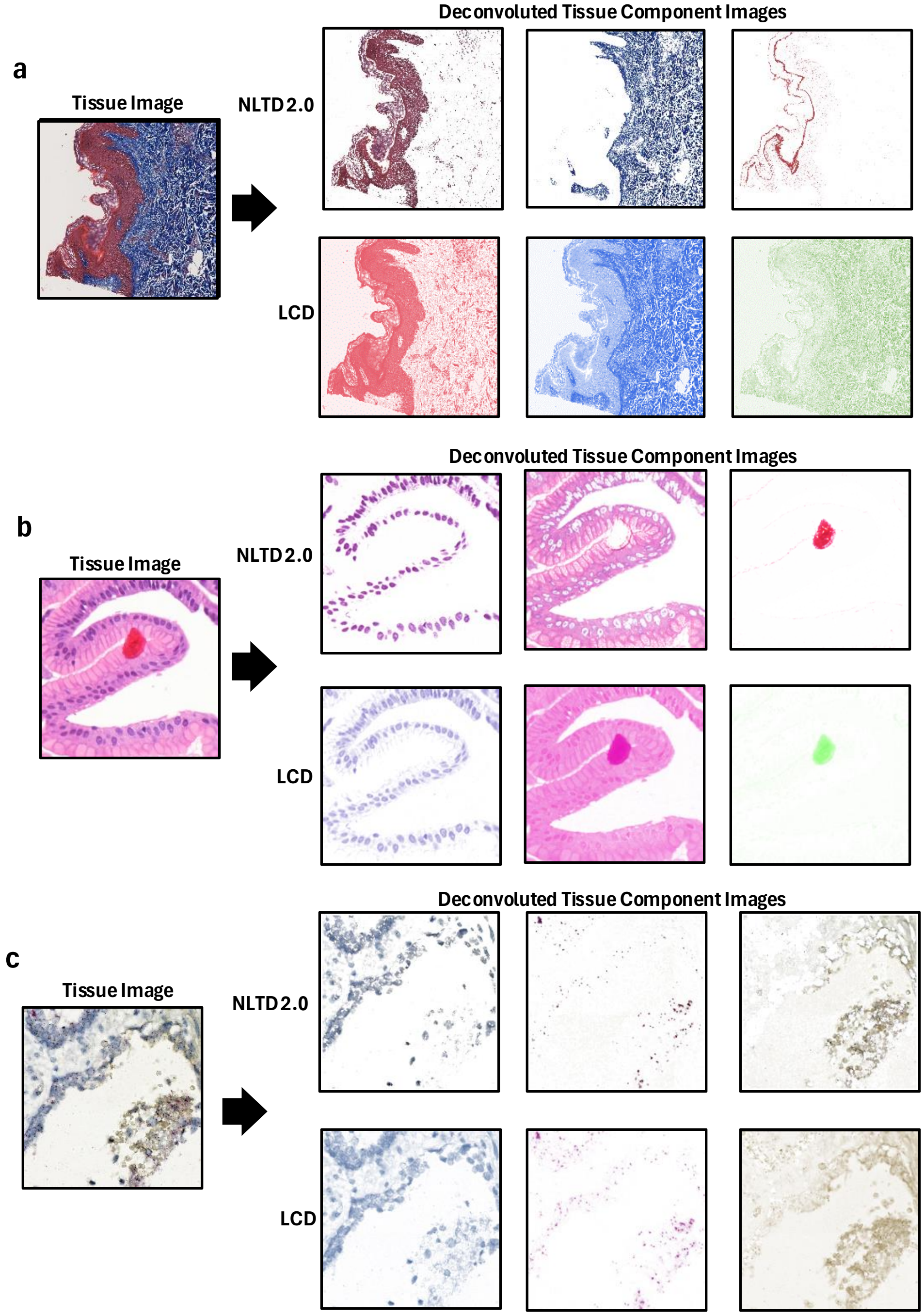


### Supplementary Fig. 3 ImageJ H&E comparison to NLTD

**a-c.** An input tissue image is shown on the left-hand side and then the performance of NLTD 2.0 and Linear Color Deconvolution (LCD) with preset vectors for each staining protocol are compared for each tissue image. In (a), a Masson's Trichrome image is used, in (b) an H&E image is used, and in (c) an IHC image is used.
